# The Genetic Architecture of Robustness for Flight Performance in *Drosophila*

**DOI:** 10.1101/2020.12.04.412395

**Authors:** Adam N. Spierer, David M. Rand

## Abstract

A central challenge of quantitative genetics is partitioning phenotypic variation into genetic and non-genetic components. These non-genetic components are usually interpreted as environmental effects; however, variation between genetically identical individuals in a common environment can still exhibit phenotypic variation. A trait’s resistance to variation is called robustness, though the genetics underlying it are poorly understood. Accordingly, we performed an association study on a previously studied, whole organism trait: flight performance. Using 197 of the Drosophila Genetic Reference Panel (DGRP) lines, we surveyed variation at the level of single nucleotide polymorphisms and whole genes using additive, marginal, and epistatic analyses that associated with robustness for flight performance. Many genes had developmental and neurodevelopmental annotations, and many more were identified from associations that differed between sexes. Additionally, many genes were pleiotropic, with several annotated for fitness-associated traits (e.g. gametogenesis and courtship). Our results corroborate a previous study for genetic modifiers of micro-environmental variation, and have sizable overlap with studies for modifiers of wing morphology and courtship behavior. These results point to an important and shared role for genetic modifiers of robustness of flight performance affecting development, neurodevelopment, and behavior.

## INTRODUCTION

Evolution acts on the genetic variation underlying phenotypic variation among individuals and populations. While many research programs focus on understanding genetic factors that contribute to phenotypic variation, fewer focus on non-genetic factors. The phenomenon of non-genetic (micro-environmental) variation describes the phenotypic variation that occurs in the absence of genetic variation, best studied in the genetically identical individuals. Non-genetic variation can arise from external (environmental) or internal (developmental) factors. Phenotypic variation across different environmental conditions (e.g. temperature) in genetically homogenous organisms is termed phenotypic plasticity. However, significant phenotypic variation can also arise among genetically homogeneous organisms in the absence of explicit environmental variation. Here, internal factors spur developmental noise in stochastic molecular processes (e.g. important transcripts or signals in very low abundance) that can result in varying levels of developmental stability (Albayrak *et al*. 2016; Schor *et al*. 2017; Klingenberg 2019). The processes or ability for organisms to maintain a consistent phenotype in the presence of these perturbations is termed buffering, while the resulting phenotype is deemed robustness (Klingenberg 2019).

Developmental noise can affect an organism’s developmental trajectory, which may impact the efficacy of natural selection by altering the association between genotype and phenotype. While it is difficult to directly observe developmental noise, deviations from an expected phenotype provide an adequate lens for study (Morgante *et al*. 2015; Vogt 2015). An example of this is deviations in bilateral symmetry (fluctuating asymmetry) (Valen 1962; Soto *et al*. 2008), which are hypothesized to be negatively associated with fitness in the case of facial symmetry (Quinto-Sanchez *et al*. 2018; Lajus *et al*. 2019). Some genetic safeguards exist to buffer against developmental noise and maintain phenotypic robustness in the presence of these stressors. Chaperonins (HSP90) do so by maintaining a protein’s structure during stressful times (Rutherford and Lindquist 1998; Chen and Wagner 2012), as does the mitochondrial unfolded protein response in maintaining homeostasis and promoting longevity (Pellegrino *et al*. 2013; Jovaisaite *et al*. 2014). In contrast, certain neurodevelopmental cell-cell adhesion molecules (e.g. DSCAMs, cadherins, and teneurins) leverage developmental noise to create more robust neural networks. In doing so, they drive repeatable non-genetic phenotypic variation in behavioral responses to serve as a bet hedging strategy (Vogt *et al*. 2008; Ayroles *et al*. 2015; Hiesinger and Hassan 2018; Honegger and de Bivort 2018).

Genes that modulate a system’s ability to resist developmental noise or a stressor are hypothesized evolutionary targets (Wagner 2008; Vogt 2015; Menezes *et al*. 2018) and subject to natural selection. And yet, these sources of non-genetic phenotypic variation are poorly understood. Previous studies have employed a Genome Wide Association Study (GWAS) framework on trait robustness, demonstrating the strategy’s feasibility as they identify significant genetic modifiers (Kain *et al*. 2012; Ayroles *et al*. 2015; Morgante *et al*. 2015; Menezes *et al*. 2018; Roman *et al*. 2018). Similarly, we sought to elucidate these genetic factors by studying the robustness of flight performance in a GWAS framework. We turned to the Drosophila Genetics Reference Panel (DGRP) lines, a collection of 205 genetically distinct and inbred lines of *D. melanogaster* that represent a snapshot of natural variation in a wild population (Mackay *et al*. 2012; Huang *et al*. 2014). Using a flight column to test flies’ ability to react and respond to an abrupt drop (Benzer 1973; Babcock and Ganetzky 2014), we tested 197 DGRP lines for their mean-normalized standard deviation (coefficient of variation) in flight performance. The coefficient of variation serves as a proxy for studying phenotypic robustness for genetically distinct groups comprised of genetically identical individuals. We identified significant additive, marginal, and epistatic variants, as well as whole gene effects on flight performance, across four sex-based phenotypes (males, females, and the average (sex-average) and difference (sex-difference) between sexes). Using defined insertional mutations in several candidate genes (*bru1, CadN, CG15236, CG32181/Adgf-A/Adgf-A2, CG3222, flippy*/*CG9766, CREG, Dscam4, flapper*/*CG11073, Form3, fry, Lasp/CG9692, Pde6, Snoo*), we successfully validated their gene roles affecting robustness of flight performance. These gene annotations suggest important roles for general and neurodevelopmental processes, and regulation of gene expression. Additionally, we found important roles for pleiotropic genes and fitness-associated genes, suggesting potential sources of developmental noise that may affect developmental stability. Together, these results contribute to a growing body of literature surrounding the genetics of robustness and corroborate previous screens for micro-environmental plasticity (Morgante *et al*. 2015), wing morphology (Pitchers *et al*. 2019), and courtship behaviors (Turner *et al*. 2013; Gaertner *et al*. 2015).

## METHODS

### Drosophila Stocks and Husbandry

197 Drosophila Genetic Reference Panel (DGRP) lines (Huang *et al*. 2014) and 24 stocks used in the validation experiment were obtained from Bloomington Drosophila Stock Center (Table S1; https://bdsc.indiana.edu/). Flies were grown on a standard cornmeal media (Mossman *et al*. 2016) at 25° under a 12h:12h light-dark cycle. Two to three days post-eclosion, they were sorted by sex under light CO2 anesthesia and given five days to recover before assaying flight performance.

### Flight performance assay

We tested approximately 100 flies of each sex from 197 DGRP genotypes (Table S1) using a refined protocol (Babcock and Ganetzky 2014) for measuring flight performance (Benzer 1973). For each sex-genotype combination, groups of 20 flies in five glass vials were knocked down, uncorked, and rapidly inverted down a 25 cm chute. The vials traveled until they reached a stop, at which point flies were ejected into a 100 cm long by 13.5 cm wide tube. Freefalling flies instinctively attempt to right themselves and land. A transparent acrylic sheet coated in TangleTrap adhesive lined the inside of the tube and immobilized flies at their respective landing height. The sheet, was removed, pinned to a white poster board, and photographed using a Raspberry Pi (model 3 B+) and PiCamera (V2). The positional coordinates were extracted using ImageJ/FIJI’s ‘Find Maxima’ feature with options for a light background and noise tolerance of 30 (Schindelin *et al*. 2012). The distributions of landing heights for each sex-genotype combination were used to calculate the mean and standard deviation. The coefficient of variation, or the mean-normalized standard deviation, was used as the final phenotype value to represent robustness.

### Genome wide association mapping

Robustness phenotypes (Table S2) were submitted to the DGRP2 webserver for the additive association analysis (http://dgrp2.gnets.ncsu.edu/) (Mackay *et al*. 2012; Huang *et al*. 2014), which returned association results for four sex-based phenotypes: males, females, average between sexes (sex-average) and difference between sexes (sex-difference). We analyzed 1,901,174 common variants (minor allele frequency ≥ 0.05) using a mixed effect model to account for Wolbachia infection status and presence of five major inversions. Since certain inversions covaried with the robustness phenotype (Table S3), only significance scores from a linear mixed model accounting for Wolbachia status and the presence of five major inversions were considered.

### Validating candidate genes

Candidate genes (Table S1B) were selected if they were identified from variants identified in the sex-average additive variant screen for mean landing height and if there were publicly available lines containing a *Minos* enhancer trap (*Mi{ET1}*) mutational insertion (Metaxakis *et al*. 2005) generated by the Drosophila Gene Disruption Project (Bellen *et al*. 2011). Experimental and control lines were derived from common isoparental crosses for each candidate gene stock backcrossed for five generations to the respective w^1118^ or y^1^w^67c23^ background. Isoparental crosses between the resulting heterozygous offspring were partitioned for absence (control line) or presence (experimental line) of the *Mi{ET1}* construct. Experimental lines were verified for homozygosity if all progeny contained the insertion after several rounds of culturing. Validations were conducting in the flight performance assay described above. The distributions in landing heights were assessed for significance if they passed a *P* ≤ 0.05 significance threshold in a Kolmogorov-Smirnov test comparing control and mutant genotypes.

### Calculating gene-score significance

Gene-level significance scores (gene-score) were determined using `PEGASUS_flies` (Spierer *et al*. 2020), a Drosophila-optimized method for the human-based platform Precise, Efficient Gene Association Score Using SNPs (`PEGASUS`) (Nakka *et al*. 2016). This analysis calculates gene-scores for each gene as a test of whether the distribution of additive variants within a gene (accounting for linkage disequilibrium) deviates from a null chi-squared distribution. Variants from the additive association screen were considered and mapped onto gene annotations and linkage disequilibrium files available with the `PEGASUS_flies` package—derived initially from the DGRP2 webserver.

### Screening for epistatic interactions

Marginal variants, corresponding with variants more likely to interact with other variants, were identified using MArginal ePIstasis Test (`MAPIT`) (Crawford *et al*. 2017). This approach uses a Bayesian framework to test the marginal effect of each variant against a focal phenotype. `MAPIT` requires a complete genotype-phenotype matrix so the DGRP genome was imputed for missing variants using BEAGLE 4.1 (Browning and Browning 2007; Browning and Browning 2016) and filtered for MAF ≥ 0.05 using VCFtools (v.0.1.16) (Danecek *et al*. 2011).

`MAPIT` was run using the ‘Davies’ method on the DGRP2 webserver’s adjusted phenotype scores, 1,952,233 imputed and filtered variants, and relatedness and covariate status files available on the DGRP2 webserver (http://dgrp2.gnets.ncsu.edu/data.html). Marginal effect P-values (data available File S8) for each sex-based phenotype were filtered for a Bonferroni threshold (P < 2.56e-8) and served as a focused subset for targeted pairwise epistasis testing against the unimputed variants (n = 1,901,174). Epistatic interactions were calculated using the ‘– epistasis’ test in a ‘–set-by-all’ framework in `PLINK` (v.1.90) (Purcell *et al*. 2007).

Significant epistatic interactions were considered if they passed a Bonferroni threshold: 0.05 / (n x 1901174 variants), where ‘n’ represents the number of significant marginal variants tested in a sex-specific subset (Table 3).

### Annotating FBgn and orthologs

FB5.57 annotations for FlyBase gene numbers (FBgn) were converted to FB_2020_01 annotations using the FlyBase tool ‘Upload/Convert IDs’ (Thurmond *et al*. 2019).

Updated FBgn (Dmel) were mapped to human orthologs (Hsap) using the Drosophila RNAi Stock Center (DRSC) Integrative Ortholog Prediction Tool (DIOPT)(Hu *et al*. 2011) tool, with the additional filtering parameter: *“Return only best match when there is more than one match per input gene or protein.”* Annotations for various genes without a citation were done so with auto-generated summaries and unreferenced descriptors of gene functions, expression profiles, and orthologs from FlyBase (Grumbling *et al*. 2006; dos Santos *et al*. 2015). These descriptors were compiled from data supplied by the Gene Ontology Consortium (Ashburner *et al*. 2000; Carbon *et al*. 2019), the Berkeley Drosophila Genome Project(Frise *et al*. 2010), FlyAtlas (Chintapalli *et al*. 2007), The Alliance of Genome Resources Consortium (Consortium 2020), modENCODE (dos Santos *et al*. 2015), DRSC Integrative Ortholog Prediction Tool (DIOPT) (Hu *et al*. 2011), and Phylogenetic Annotation and INference Tool (PAINT) (Gaudet *et al*. 2011).

### Data availability

All phenotype data required to run the outlined analyses are available in Table S2 or using the DGRP2 webserver (http://dgrp2.gnets.ncsu.edu/). All supporting information is available at: https://doi.org/10.7910/DVN/RSJNW5.

## RESULTS and DISCUSSION

We sought to identify the genetic modifiers of robustness in a whole organism phenotype: flight performance. Using the Drosophila Genetic Reference Panel (DGRP) lines, we identified several additive, marginal, and epistatic variants, as well as whole genes that associate with genotype robustness in response to a flight challenge. In the sections that follow we describe the variant-based, gene-based, and epistatic analyses in turn.

### Variation in flight performance across the DGRP

We screened 197 DGRP lines (Table S1) for their flight ability in response to an abrupt drop (Figure 1A). Qualitative observations made in a previous study of strong, intermediate, and weak genotypes in the flight assay suggests stronger genotypes react faster and respond more effectively than weaker ones (Spierer *et al*. 2020). The mean and standard deviation in landing height were calculated for each sex-genotype combination, along with the mean-normalized standard deviation (coefficient of variation). The coefficient of variation served as our metric for robustness (Figure S1; Table S3). Genotypes with a lower coefficient of variation (more consistent) are more robust for flight performance (Klingenberg 2019). On average, flight performance was more robust in males than females (males: 0.17 A.U. ± 0.055 SD vs. females: 0.22 A.U. ± 0.075 SD; Figures 1B and S2). There was a significant relationship in robustness between sexes (R = 0.55; *P* < 1E-16; Figure 1C), suggesting the genetic architecture of robustness in flight performance is similar between the sexes. However, the magnitude of the regression coefficient suggests robustness is somewhat sexual dimorphic.

**Figure 1.**
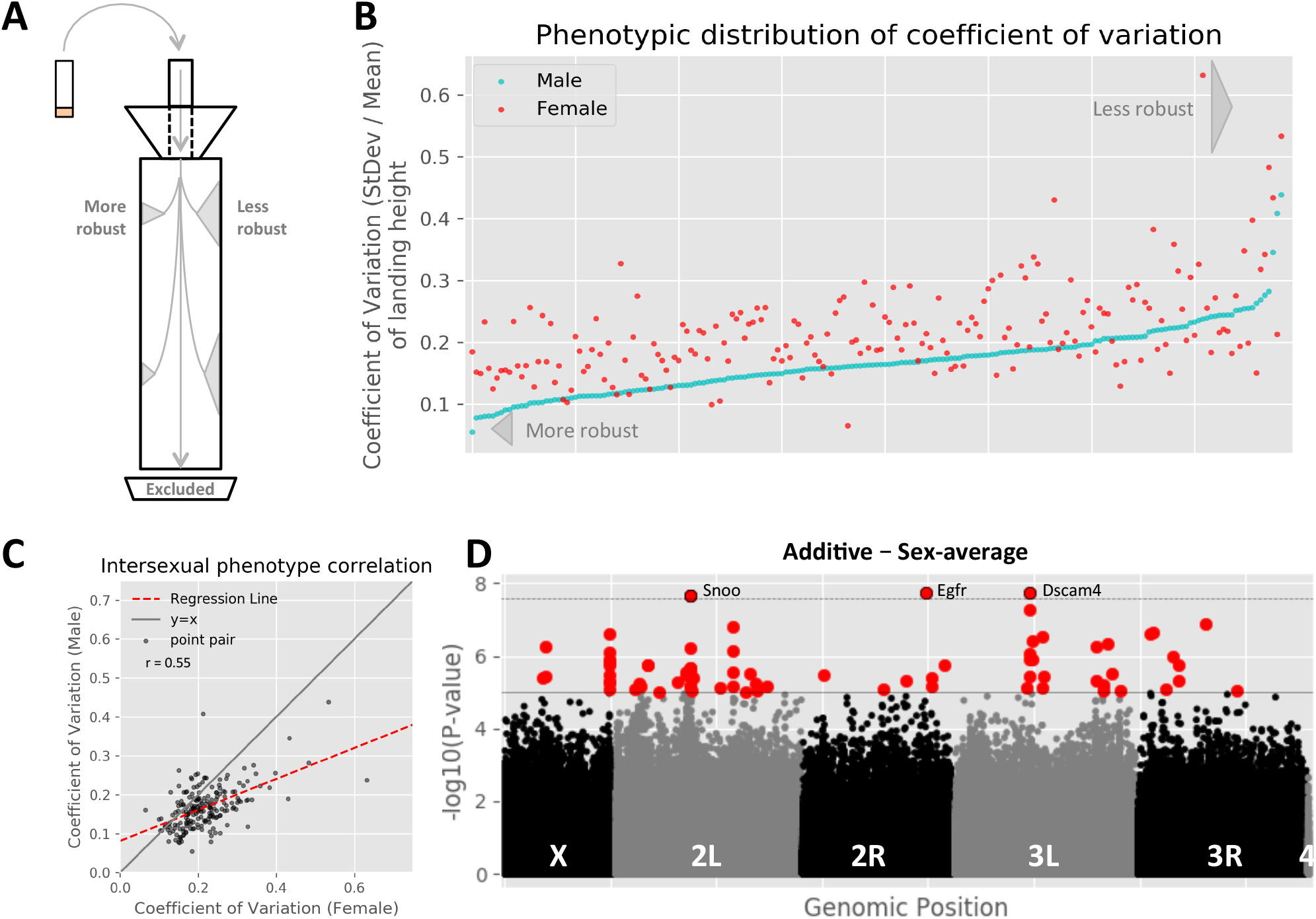
The *Drosophila* Genetic Reference Panel lines demonstrate variation for robustness in flight performance across genotypes and sexes. (A) Flies were assayed for flight performance using a meter-long flight column (Babcock and Ganetzky 2014). The coefficient of variation (mean-normalized standard deviation) is a proxy for robustness; more robust genotypes have less variation in landing height around the mean. Flies that passed through the column were excluded from the analysis. (B) The phenotypic distribution of sex-genotype pairs, ordered by increasing male score, demonstrates the DGRP lines have variation in their robustness for flight performance. Genotypes demonstrated phenotypic variation for robustness in both sexes. (C) Males were generally more robust than females, though the two were related (R = 0.55, *P* < 1E-16; regression line in red). Sexual dimorphism is observed by the intersection of the regression line and y = x line (gray). (D) Additive variants in the sex-average analysis, visualized as a function of the –log10 of variants’ *P*-value illustrates several variants (red) passed the traditional DGRP significance threshold (*P* ≤ 1E-5; gray solid line), and three (red with black outline) passed Bonferroni significance threshold (*P* ≤ 2.63E-8, gray dashed line). Variants that did not pass the significance threshold are colored in black or gray by chromosome.

We tested our phenotype in each sex against those publicly available on the DGRP2 webserver to determine whether robustness of flight performance was a unique trait. Indeed, we found no significant relationship after applying a multiple testing correction (*P* ≤ 1.85E-3; Table S3), suggesting it is unique. Accordingly, we took four distinct approaches to find significant associations from different facets of the genetic architecture (Table 1).

**Table 1.** Different approaches uncover different types of genetic modifiers affecting the focal phenotype. No single screen will identify all modifiers, so four overlapping approaches were conducted to better survey the genetic architecture of robustness of flight performance.

|  |  |  |
| --- | --- | --- |
| **Screen type** | **Target modifier type** | **Analysis platform** |
| Additive | Variants of large effect | DGRP2 webserver/FastLMM |
| Marginal | Interconnected variants | MAPIT |
| Epistatic interactions | Connections between variants | PLINK |
| Whole gene | Genes of moderate effect | PEGASUS_flies |

### Several variants of large effect associate with robustness in flight performance

We performed a Genome Wide Association Study (GWAS) to calculate the significance of variants’ additive effects, and subsequently whole gene significance scores. We analyzed the effects of 1,901,174 common variants (MAF ≥ 0.05) across for four sex-based phenotypes (males, females, the sex-average, and sex-difference; Figures 1D and S3-5). Two of the major inversions covaried with our phenotype scores (Table S4), so we used a mixed model to account for Wolbachia infection status, presence of inversions, and polygenic relatedness.

We performed a traditional, additive variant screen using the DGRP2 webserver pipeline. Eight variants in the male, female, or sex-average analyses passed a strict Bonferroni threshold (*P* ≤ 2.63E-8; Tables 2-4; Table S5). Three variants (2R_17433667_SNP, 3R_4379159_SNP, 3R_9684126_SNP) were also significant for the mean landing height in flight performance (Spierer *et al*. 2020) and mapped to *Epidermal Growth Factor Receptor* (*Egfr*; human homolog *EGFR*), *Odorant receptor 85d* (*Or85d*) and an intergenic region on chromosome 3R, respectively. *Egfr* is a pleiotropic, tyrosine kinase receptor involved in several developmental and homeostatic processes, as well as playing a known source of natural variants that modify wing shape and affect flight performance (Paul *et al*. 2013; Pitchers *et al*. 2019). *Or85d* is an odorant receptor expressed on the antennae and maxillary palp (Couto *et al*. 2005). The significant variant in *Or85d* coded for a non-synonymous SNP (C277Y) at highly conserved site (Figure S6) (Siepel *et al*. 2005; Siepel and Haussler 2005), though follow up analysis with the PROVEAN webtool (Choi and Chan 2015) suggests this mutation is neutral (scored -2.312 with -2.5 as deleterious). Finally, the intergenic region lacked any embryonic transcription factor binding site (TFBS) annotations (Moses *et al*. 2006; Li *et al*. 2008; MacArthur *et al*. 2009; Thomas *et al*. 2011), suggesting it may interact with transcription factors or epigenetic factors later during development or homeostasis.

**Table 2.** Eight additive variants passed the Bonferroni threshold. In the additive approach, eight variants passed the strict Bonferroni significance threshold (*P* ≤ 2.63E-8). These common variants were typically near the Minor Allele Frequency (MAF) threshold of 0.05. Nearly all variants mapped to genes, three of which had human homologs. Non-coding variants mapped to introns or upstream of the gene’s coding region, however three variants also contained transcription factor binding sites (TFBS) annotated active during the embryonic stage (NEGRE *et al*. 2011).

| Variant | MAF | Variant type | Annotation |  |  |
| --- | --- | --- | --- | --- | --- |
|  |  |  | Gene symbol |  | Embryonic TFBS |
|  |  |  | Dmel | Hsap |  |
| 2L_7949902_SNP | 0.053 | Intron | *Snoo* | *SKI* |  |
| 2L_7949906_SNP | 0.053 | Intron | *Snoo* | *SKI* | - |
| 2R_17433667_SNP | 0.051 | Intron | *Egfr* | *EGFR* | *bcd, da, dl, gt, hb, kni, Med, prd, sna, tll, twi, disco, Trl* |
| 2R_17987191_SNP | 0.067 | Upstream (152 bp) | *Flapper/CG11073* | - | *dl* |
| 2R_17987203_SNP | 0.062 | Upstream (164 bp) | *Flapper/CG11073* | - | *dl* |
| 3L_8237797_DEL | 0.084 | Intron | *Dscam4* | *DSCAM* | - |
| 3R_4379159_SNP | 0.053 | Non-synonymous | *Or85d* | - | - |
| 3R_9684126_SNP | 0.15 | Intergenic |  | - | - |

**Table 3.**
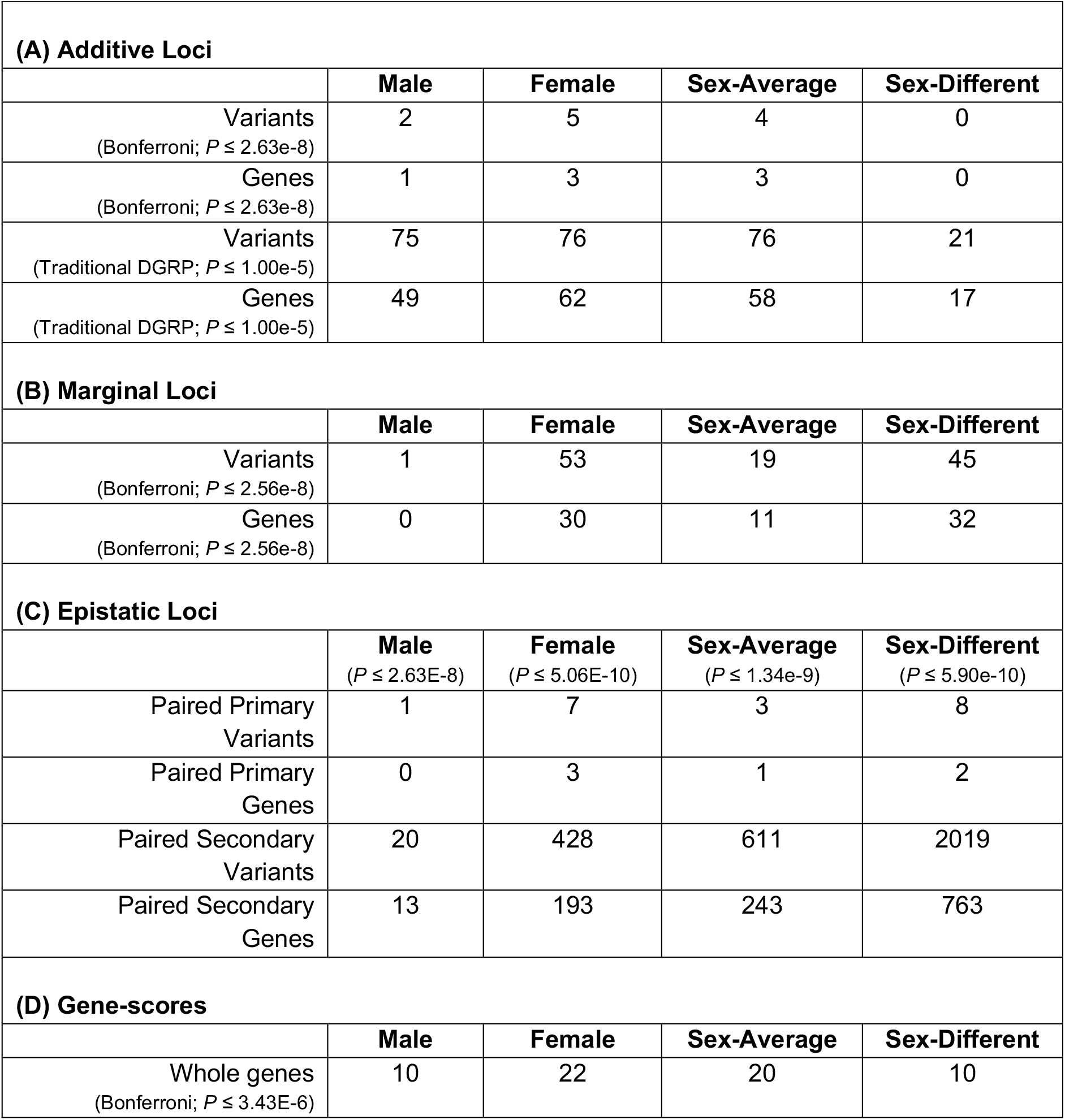
Each analysis and sex-based phenotype identified varying enrichment for genetic modifiers. (A) Additive loci (variants and the genes they map to) at Bonferroni and traditional DGRP GWAS thresholds differ in enrichment by an order of magnitude. (B) Marginal variants mapped to several genes, and were tested for (C) epistatic interactions. Marginal variants were only tested for epistatic interactions if they passed MAF ≥ 0.05 in the unimputed genome. Finally, (D) whole genes were identified consistently across all sex-based phenotypes

The remaining Bonferroni additive variants mapped to genes that were also identified from additive Bonferroni variants in the mean flight performance screen (*Dscam4* and *CG11073*/*flapper*) or were otherwise strongly significant (*Snoo*). These three genes have known or hypothesized roles in developing robust neural circuits (Quijano *et al*. 2010; Tadros *et al*. 2016; Spierer *et al*. 2020). Their identification in mean and robustness of flight performance screens suggests they have a dual role in affecting a genotype’s ability and variability in flight performance.

Applying a traditional DGRP association threshold (*P* ≤ 1E-5), we observed a trend in our data where neural and pleiotropic genes play an important role in the genetic architecture of robustness of flight performance. We identified 163 unique, significant variants (Table S5), 18 of which mapped to coding regions (Table 4). These coding variants include a novel transcriptional start site in a gene (*CG43707*) affecting muscle architecture and flight performance (Schnorrer *et al*. 2010) and six non-synonymous SNPs that each mapped to unique genes (*CG12517, CG13794, CG34215, Or85d, Spn, Tif-IA*). Some of these affect neural phenotypes, like *CG13794*, a neurotransmitter (Couto *et al*. 2005; Romero-Calderon *et al*. 2007), while others are pleiotropic and affect multiple traits. *CG12517* and *Tif-IA* are involved in the stress response of the fat body and insulin-based metabolism, respectively, and both are involved in development of the germline (Yatsu *et al*. 2008; Tootle *et al*. 2011; Tsuzuki *et al*. 2012; Ghosh *et al*. 2014). *Spn (Spinophilin*; human homolog *PPP1R9A*), a pre-synaptic regulator of neurons (Muhammad *et al*. 2015), affects flight performance (Schnorrer *et al*. 2010), male aggression (Edwards *et al*. 2009), odor response (Sambandan *et al*. 2006), and is also found in sperm (Wasbrough *et al*. 2010).

**Table 4.**
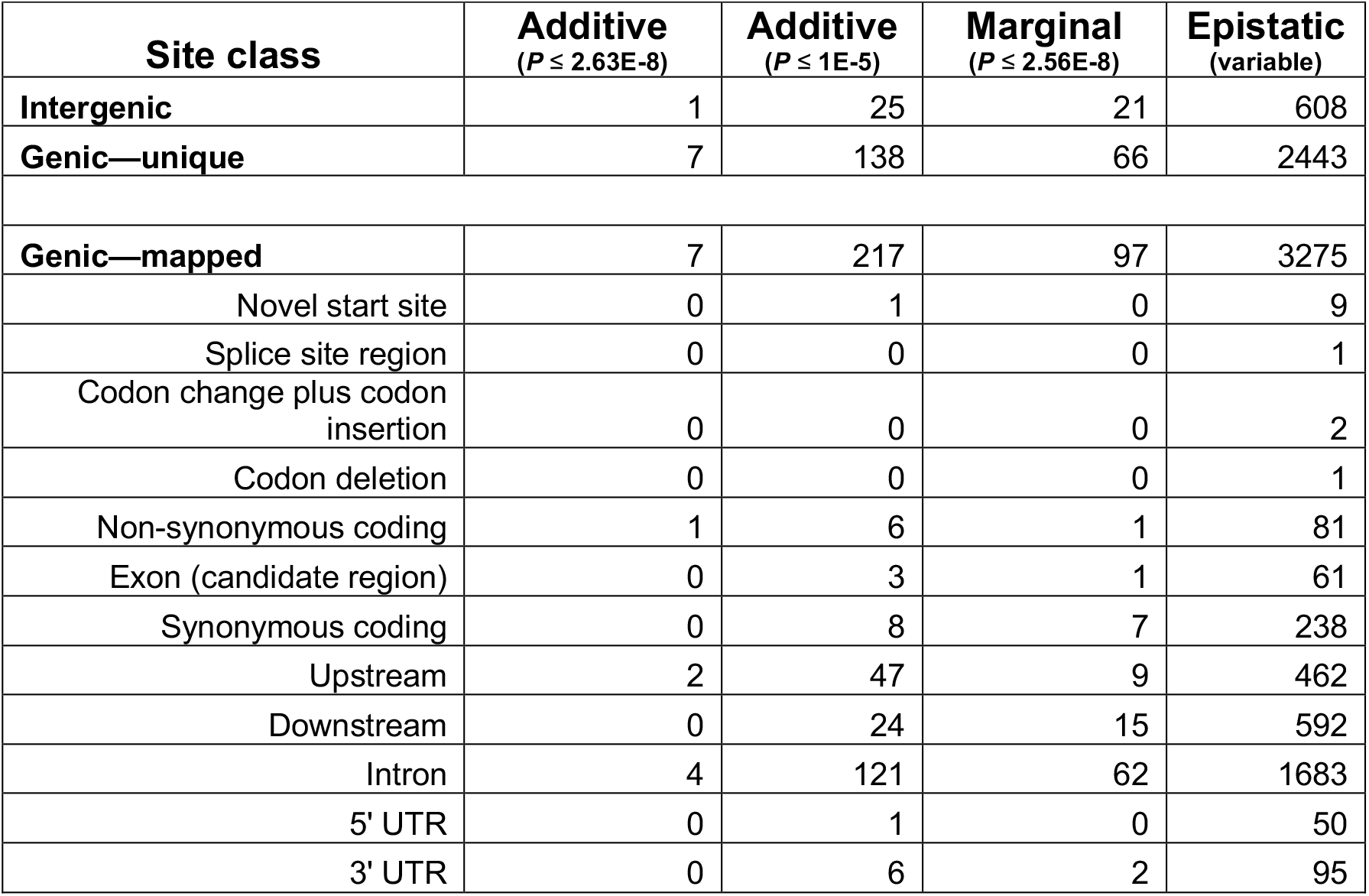
Significant variants are non-uniformly distributed across site classes. Variants mapped to several site classes in the genome. Across all variant-based analyses, intergenic (variants lacking FBgn annotations) and genic (sites with FBgn annotations) sites were both represented. Some genic variants mapped to multiple FBgn (Genic—mapped). These genic variants mapped to 12 different site classes.

|  |  |  |  |  |
| --- | --- | --- | --- | --- |
| Site class | Additive<br>($P \leq 2.63E-8$) | Additive<br>($P \leq 1E-5$) | Marginal<br>($P \leq 2.56E-8$) | Epistatic<br>(variable) |
| Intergenic | 1 | 25 | 21 | 608 |
| Genic—unique | 7 | 138 | 66 | 2443 |
| Genic—mapped | 7 | 217 | 97 | 3275 |
| Novel start site | 0 | 1 | 0 | 9 |
| Splice site region | 0 | 0 | 0 | 1 |
| Codon change plus codon insertion | 0 | 0 | 0 | 2 |
| Codon deletion | 0 | 0 | 0 | 1 |
| Non-synonymous coding | 1 | 6 | 1 | 81 |
| Exon (candidate region) | 0 | 3 | 1 | 61 |
| Synonymous coding | 0 | 8 | 7 | 238 |
| Upstream | 2 | 47 | 9 | 462 |
| Downstream | 0 | 24 | 15 | 592 |
| Intron | 4 | 121 | 62 | 1683 |
| 5' UTR | 0 | 1 | 0 | 50 |
| 3' UTR | 0 | 6 | 2 | 95 |

Most variants mapped to complex traits with the DGRP are enriched for intergenic and non-coding regions (Table 4) and have presumably regulatory roles (Mackay *et al*. 2012; Mackay and Huang 2018). Non-coding variants mapped to genes with important roles in development and function of the nervous system, wing morphology, muscle function, and regulation of gene expression. They also play important roles in gametogenesis or determination of sex identity, establishing a possible connection between genotype and the sexually dimorphic phenotype we measured. A table of all genes found in the current study is available as a master lookup table (File S1). The identification of these genes and their respective annotations may hint at additional genetic sources of developmental stabilization.

### Functional validation of candidate genes supports a role for neurodevelopment affecting robustness of flight performance

We functionally validated several genes’ roles in affecting robustness of flight performance. Using insertional mutants in the candidate genes identified from the mean landing height screen, we applied a Kolmogrov-Smirnov test between insert and backcross-control genotypes (see methods) to test for differences in the distribution of landing heights. We validated 11 single genes (*bru1, CadN, flippy (CG9766), CG15236, CREG, Dscam4, flapper (CG11073), form3, fry, Pde6*, and *Snoo*), and two constructs that fell in multiple genes (*Adgf-A/Adgf-A2/CG32181 and CG9692/Lasp*) (Figures 2 and S7; Table S6). These genes are putative sources of genetic variation in maintaining developmental stability. They were also validated in the mean flight performance screen (Spierer *et al*. 2020), suggesting these genes likely play dual roles modifying the ability and variability of flight performance.

**Figure 2.**
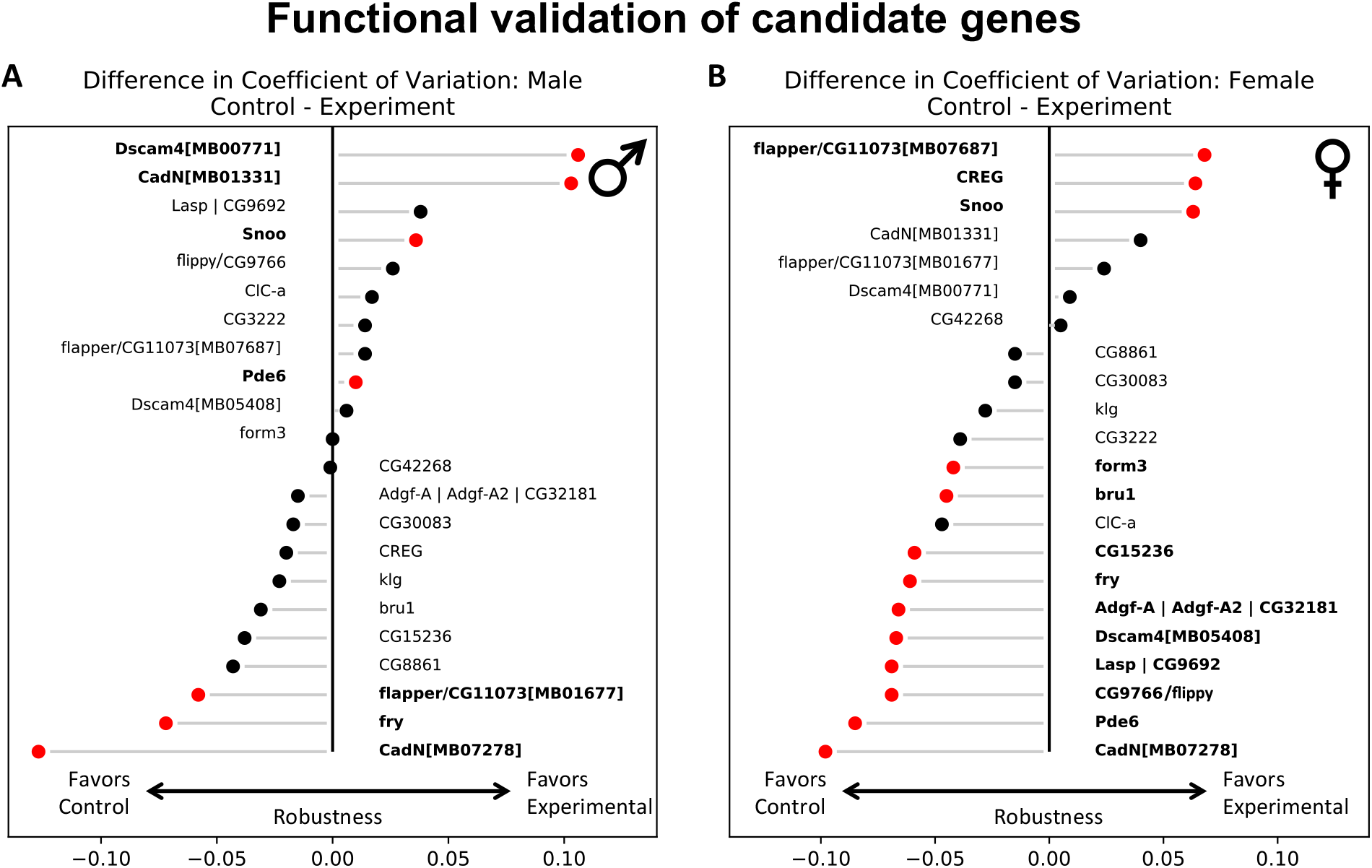
Several genes validated for robustness of flight performance. Flies homozygous for *Mi{ET1}* insertion constructs inserted in candidate genes (experiment) were tested against their background control (control). Comparisons between control and experiment lines were assessed for significance using a Kolmogrov-Smirnoff test (*P* ≤ 0.05; red points and bold text). Values to the left of the midline suggest control genotypes were more robust than experimental lines, while the opposite is true for values to the right of the line. (A) Seven constructs were significant in males, (B) while 13 were significant in females. Some candidate genes were tested more than once (*CadN, Dscam4*, and *flapper*) because they were strongly significant in the sex-average additive association screen. Separate constructs are denoted by a suffix containing a ‘MB’ code.

### Analyses of whole-gene effects identifies distinct factors affecting robustness

The additive variant screen takes a conventional minSNP approach, deeming a gene significant if its most significant variant passes a significance threshold. However, this approach is biased toward longer genes and does not account for linkage between sites. To counteract these biases, we employed `PEGASUS_flies` (Spierer *et al*. 2020), a *Drosophila* version of the human-focused `PEGASUS` platform (Nakka *et al*. 2016). This method takes a gene-specific approach; assessing a whole gene’s significance by testing the distribution of variants within a gene against a null chi-squared distribution of SNP *P*-values. Thus, it can detect significant genes of moderate effect, as well as genes that may be missed in a minSNP approach.

Using `PEGASUS_flies`, we identified 45 unique genes (Table S7) across all four sex-based phenotypes that passed a Bonferroni threshold (*P* ≤ 3.43E-6; Figures 3A and S8). Two were present in the additive screen (*nmo* and *Sdc*), along with 27 other genes with annotations for neurodevelopment and function (*ana3, barc, Br140, caps, CG5921, CG5937, CG12163, CG44774, Crz, ct, ctrip, Dop2R, Dys, ena, ham, Nckx30C, Oct-TyR, olf186-F, PsGEF, Ptp4E, rad, rodgi, row, tou, TTLL5, tutl, wde*). Some genes also affected muscle, chaete, or general development (*caps, CG5937, CG31635, CG32521, CG3277, CG43333)*, while others facilitated gametogenesis or promoted reproductive success (*ana3, CG1632, CG5937, CG12163, CG44774, CHES-1-like, Crz, ct, Dop2R, Dys, ena, Gbs-70E, PsGEF, tou, wde*). These results largely corroborate annotations from the additive search and expanded the number of genes that associate with robustness in flight performance.

**Figure 3.**
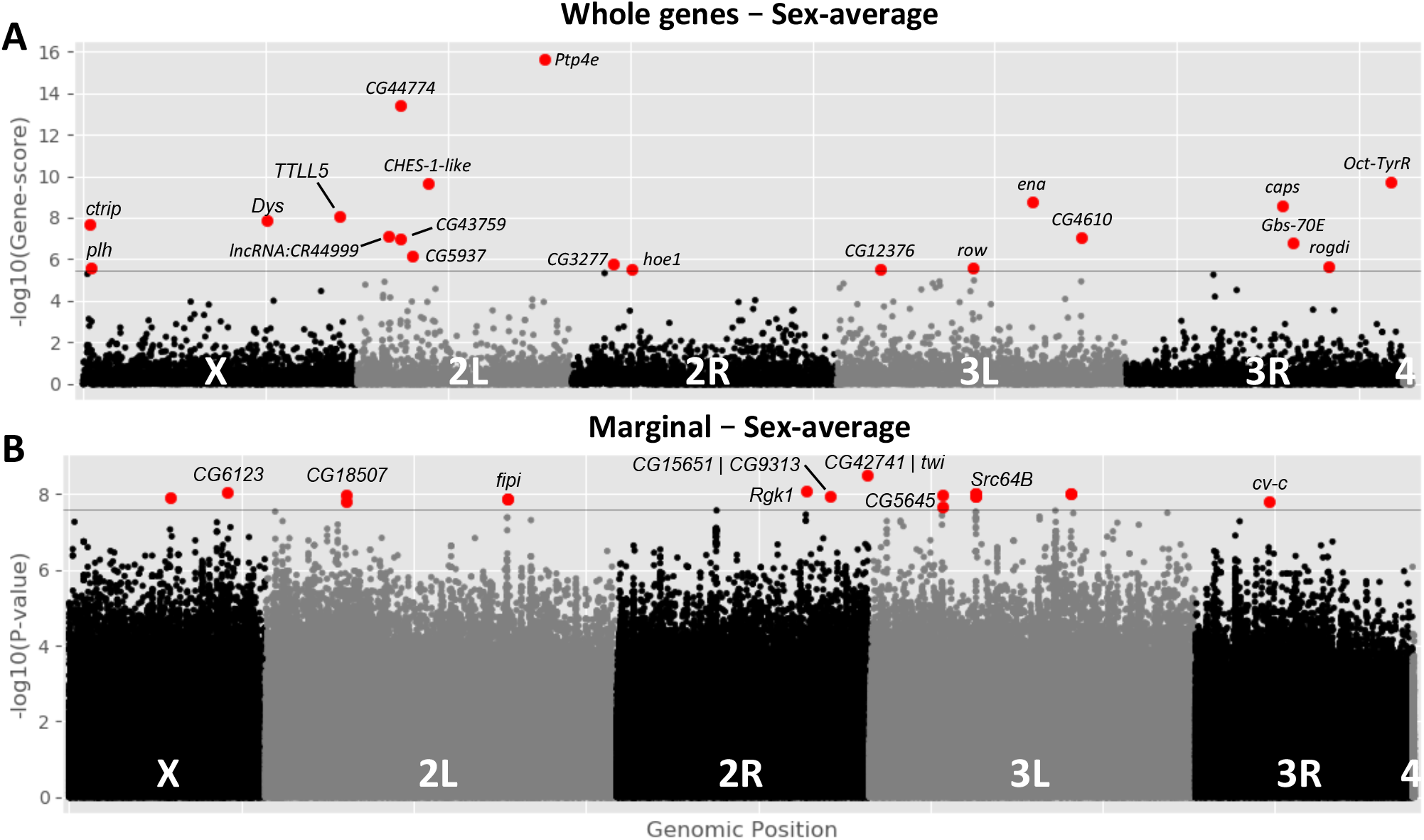
Several genetic variants positively associate with flight performance across different types of analyses. (A) Manhattan plot for sex-average whole gene analysis suggests several genes (red) were significant above a Bonferroni threshold (*P* ≤ 3.43E-6, gray line). (B) Manhattan plot for sex-average marginal analysis suggests several variants (red) were significant above a Bonferroni threshold (*P* ≤ 2.56E-8, gray line). For each plot, points are arranged by relative chromosome (genomic) position and all points are –log10 transformed.

### Association of marginal and epistatic variants with robustness in flight performance

Epistatic, or pairwise, interactions play an outsized role as context-specific effectors in complex traits (Huang *et al*. 2012). Traditional epistasis analyses face large computational and statistical hurdles, so we turned to MArginal ePIstasis Test (`MAPIT`) to focus the search for interactions (Crawford *et al*. 2017). `MAPIT` is a Bayesian approach that identifies marginal variants representing genetic hubs. Using this subset of hub variants, we performed a set-by-all epistasis search against all other variants.

We identified 104 significant marginal variants in 66 genes and several intergenic regions that exceeded a Bonferroni threshold (*P* ≤ 2.56E-8) across all sex-based phenotypes (Figures 3B and S9; Table S8). Most of these variants mapped to non-coding regions, suggesting an important regulatory role among marginal variants (Mackay and Huang 2018). Among the 19 the significant marginal variants with epistatic interactions (discussed below), only seven mapped to genes (*Bx, CG9313, CG15651, CG9171, PKC-δ/Pkcdelta, jvl, ush*).

In total, there were 6313 significant epistatic interactions that mapped to 1081 genes (Table 3). Interestingly, several of the marginal genes (containing marginal variants) had epistatic interactions with other marginal genes (marginal-marginal epistatic interactions; Figure 4A). However, these interactions were typically between a marginal and non-marginal variant, rather than between two marginal variants. This finding suggests a highly interconnected genetic architecture. Broadly, epistatic interactions were enriched for neurodevelopment and general development, though there were too many significant interactions to comprehensively describe in text (Table S9). Instead, we will focus on the marginal variants that mapped to genes and some of their noteworthy epistatic interactions.

**Figure 4.**
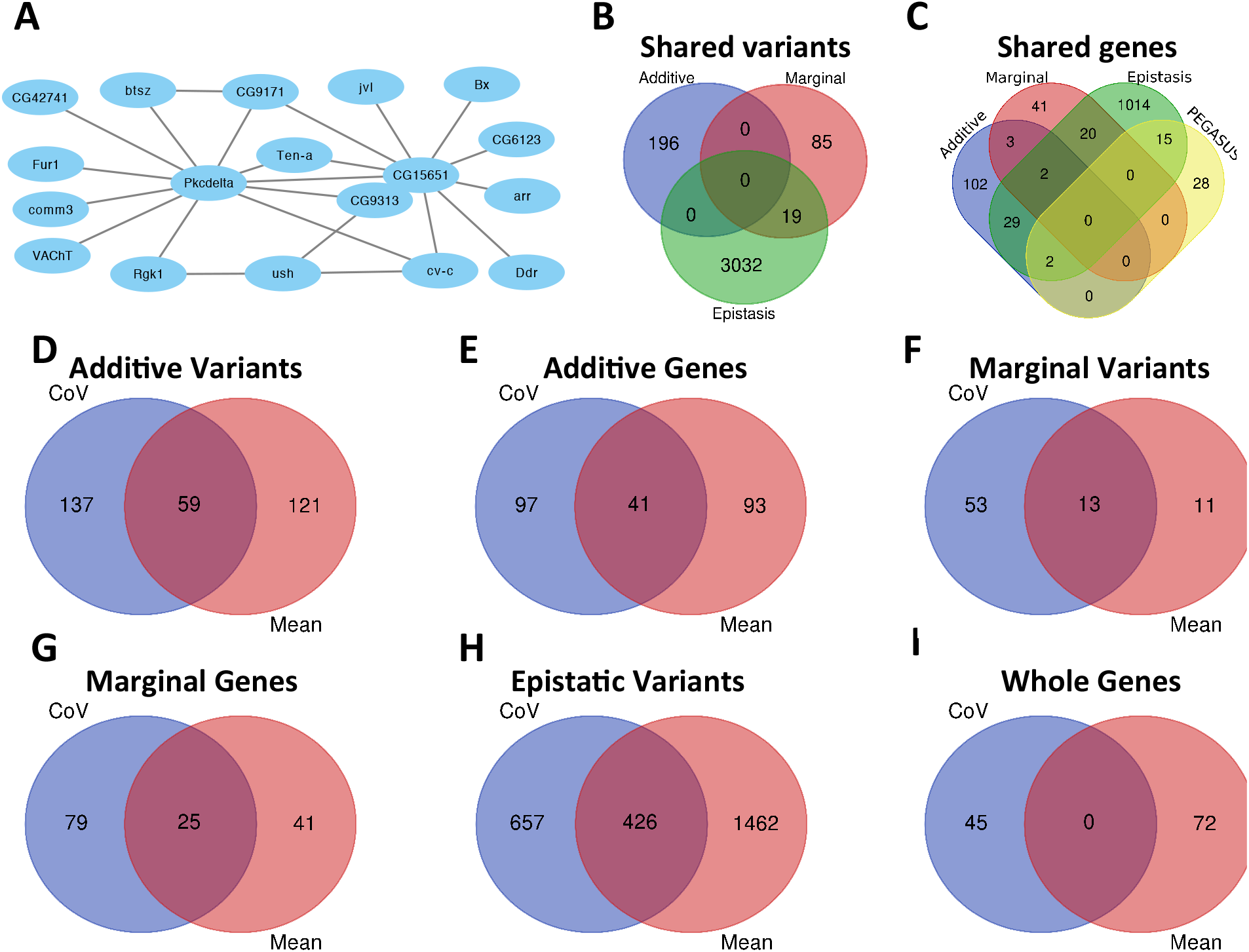
Robustness of flight performance is comprised of an interconnected genetic architecture. (A) There were several interactions between genes identified from marginal variants. In particular, *PKC-δ* had the greatest number of interactions with other marginal genes, while *CG15651* and *CG9313* were next. There was a marginal variant that overlapped with *CG15651* and *CG9313*, so all edges connecting to *CG15651* also connect to *CG9313*, however there was an independent variant in *CG9313* that did not overlap with *CG15651* that interacted in the sex-difference screen with *PCK-δ* and *ush*. Intergenic regions that also interacted with genic marginal variants are not displayed. (B) For the additive, marginal, and epistatic variants identified, additive variants were unique, while marginal and epistatic variants had some overlap. This overlap was expected since the marginal variants served as a subset in searching for epistatic variants. (C) Genes and genes mapped from minSNP-variants had some overlap between analyses, though most genes were unique to a single analysis. When comparing variants and unique genes across (D-E) additive, marginal (F-G), and epistatic (H) analyses, there was roughly 15-20% overlap between the shared group and all those identified. However, there was no overlap between the (I) whole genes identified using `PEGASUS_flies`.

### Many marginal variants in female and sex-average epistasis analyses map to pleiotropic genes

While the male marginal variant mapped to an intergenic region, there were several marginal variants in the female and sex-average analyses that mapped to genes, especially pleiotropic genes. Among these were *Beadex* (*Bx*; human homolog *LMO1*), *CG9171* (human homolog *B4GAT1*), *CG15651* (human homolog FKRP), *CG9313, and javelin-like* (*jvl*). The corresponding and significant epistatic interactions, like these marginal genes, have annotations for wing morphology, muscle development, neural circuit assembly and neuronal function, and interestingly, sex-related behaviors and sex-specific tissues (Table S8). For more information on marginal gene functions, please see “Many marginal variants in female and sex-average epistasis analyses map to pleiotropic genes, continued” in the Supplemental Results.

### Epistatic interactions associating with the sex-difference phenotype

As noted previously, there was relationship between male and female robustness scores (R = 0.55) that suggests a genetic basis for sexual dimorphism. We found a signature of this dimorphism in the sex-difference epistasis search, which had four times as many epistatic interactions as the next closest sex-based phenotype (females).

*Protein Kinase C-δ* (*PKC-δ* or *Pkcd*; human homologs *PRKCD* and *PRKCQ*) drove this trend and accounted for over half (3211 of 6313) of all epistatic interactions in our study. Some of these interactions were with variants in other marginal genes (Figure 4A), suggesting a more central and interconnected role within the genetic network. *PKC-δ* is a member of the Protein Kinase C family and is known to modulate flies ability to learn from their environment, especially during flight (Colomb and Brembs 2016; Getahun *et al*. 2016; Gorostiza *et al*. 2016). Flies inability to learn from proprioceptive cues corresponds increased variation in their flight path (Hesselberg and Lehmann 2009; Lehmann and Bartussek 2017). Thus, *PKC-δ* is a noteworthy candidate gene with a previously validated effect on flight robustness.

Another marginal variant from the sex-difference epistasis screen was the developmental transcription factor *u-shaped* (*ush*; human homolog *ZFPM1*), which mediates neurodevelopment and thorax development (Fromental-Ramain *et al*. 2010). It regulates *scute* (*sc*), with roles in the sex-determination pathway (Wrischnik *et al*. 2003), and the SC-A (*scute* and *achaete*) complex, which contributes to development of mechanosensory chaete and sensory organs on the wing (Skeath and Carroll 1991; Cubadda *et al*. 1997). Accordingly, *ush* is similarly a pleiotropic gene that acts as a potential source of developmental noise and stability affecting robustness of flight performance. For more examples and information of epistatic interactions between *ush* and other genes, see “Epistatic interactions associating with the sex-difference phenotype, continued” in the Supplemental Results.

### Bet hedging as a modifier of robustness

Bet hedging is an evolutionary strategy for increasing phenotypic variation in the presence of genetic homogeneity (Hiesinger and Hassan 2018; Honegger and de Bivort 2018). We identified putative bet hedging genes involved with neurodevelopment, such as: Down Syndrome Cell Adhesion Molecules (DSCAM; *Dscam3* and *Dscam4*), cadherins (*Cad87A, CadN, CadN2*), and teneurin (*Ten-a*) family genes. These genes were also significant interactors with *PKC-δ*. They contribute to differential wiring of the peripheral and central nervous systems through dendritic self-avoidance (Hong *et al*. 2012; Li *et al*. 2020) in the brain and sensory organs of the wing (Kise and Schmucker 2013; Nagai and Mizuno 2014). Interestingly, *Ten-a* was previously identified and validated in a screen for individuality in locomotor handedness (Ayroles *et al*. 2015) and we validated *CadN* and *Dscam4* in the present study.

Together, these genes and their roles in differential circuit assembly serve as validated modulators of robustness.

### Flight and courtship share morphological structures and genetic modifiers

Interestingly, we identified several pleiotropic genes with dual roles in robustness of flight performance and courtship and fitness-associated traits. One such gene from the marginal screen was *factor of interpulse interval* (*fipi*), a marginal variant with no significant epistatic interactors. This gene regulates the intervals of courtship song (Fedotov *et al*. 2018) and was previously identified in an independent screen for micro-environmental variation (Morgante *et al*. 2015). Other courtship-related studies with the DGRP shared genes with robustness in flight performance (*CG1358* and *Dif*) (Turner *et al*. 2013) and (*bru-3, CG13024, CG42784, Fur1, shot, SKIP, Ubx, wuc*) (Gaertner *et al*. 2015). We identified several other genes with annotations for sex determination, courtship behavior, and sex-specific neural patterning (*Alh, bab1, Btk29A, CASK, chinmo, dnc, dysb, fru, gom, lov, Mip, Nrg, Sh, Tbh*), as well as many genes with dual roles in gametogenesis (previously listed).

The link between pleiotropic genes association with both fitness-associated traits and robustness of flight performance may lie in the development of common structures. In flight, well-structured wings are important for generating lift (Marcus 2001) and chaete are important for proprioception (Furman and Bukharina 2008; Quijano *et al*. 2010). In courtship, wings are used for visual flagging and courtship song (Sadaf *et al*. 2015), while chaete act as chemosensors for pheromones (Thistle *et al*. 2012; Pavlou and Goodwin 2013) and mechanosensors in courtship. Additionally, the neural networks innervating chaete and connecting them to the central nervous system (CNS) are differentially patterned by cell-cell adhesion molecules (e.g. DSCAMS, cadherins, and teneurins) (Hong *et al*. 2012). We successfully validated *Dscam4* and *CadN*, two such neural patterning genes with important implications in differential detection of pheromones and courtship behaviors (Yu *et al*. 2010; Pavlou and Goodwin 2013), and flight performance (Spierer *et al*. 2020), lending support to our hypothesis.

These pleiotropic genes may be associated with robustness as a result of an evolutionary tug-of-war caused by intralocus sexual dimorphism or conflict. In other words, what is beneficial for one sex may be neutral or disadvantageous for the other. This phenomenon is observed in insects in the context of locomotor performance, courtship behavior, and fitness (Berger *et al*. 2014; Berger *et al*. 2016). In studies where male flies were allowed to genetically “win” the sex conflict and evolve, males increased locomotor activity, fitness, and variation in wing morphology, while females saw a reduction in locomotor activity and fitness (Long and Rice 2007; Abbott *et al*. 2010). Variation in wing morphology is noteworthy because it is hypothesized to be under stabilizing selection (Munoz-Munoz *et al*. 2016; Sztepanacz *et al*. 2017). It is also sensitive to factors that buffer against developmental noise and serves as a strong proxy for developmental stability (Soto *et al*. 2008; Klingenberg 2019). Therefore, flies with less within- and between-individual variation in wing morphology are thought to have better buffered genetic networks against developmental noise and stochasticity.

### Genetic architecture of robustness is comprised of different types of modifiers

Each analysis we conducted sheds light on different areas of the genetic architecture (Table 1). The additive variant analysis identified single variants with larger effects on the phenotype, while the whole gene analysis identified genes of moderate effect based on the distribution of additive variants in a gene. The marginal variant analysis identified variants more likely to interact with other variants, while the epistasis analysis identified those respective interactions. Of the variant-based analyses, all additive variants were exclusive, though the marginal variants and epistatic interactions had some overlap (Figure 4B). When mapping these variants to genes, all analyses identified genes that were shared with at least two other analyses (Figure 4C). This result suggests that genes contain different types of variants that affect separate facets of the genetic architecture. For a complete list of all genes identified in this study and which analysis they were present in, see File S1.

### Overlap between robustness and other DGRP studies

When comparing screens for variants associating with overall flight performance (mean) against robustness for flight performance (coefficient of variation), we consistently identified approximately 15-20% overlap between variants and their mapped genes (Figure 4D-H; Table S10) with the exception of the whole gene analysis (no overlap; Figure 4I). These results suggest that while certain main features of the genetic architecture are shared, the traits have largely separate genetic architectures.

Similarly, we found commonalities between robustness in flight performance and other DGRP studies conducted beyond the flight phenotype. A micro-environmental plasticity screen for startle response, resistance to starvation, and chill coma recovery shared 37 genes with robustness of flight performance (*Bsg25D, CARPB, CG17716, CG31690, CG32767, CG33981, CG4168, CG42322, CG42324, CG43901, CG5853, Diap1, dpr6, dpr8, E2f1, ed, Eip63E, FAM21, fipi, fred, fru, IA-2, Lac, Lmpt, lncRNA:CR32773, lncRNA:iab8, Moe, mtgo, nub, Pde9, PsGEF, Ptp99A, pum, Pvf3, rdgA, Rgk3, Src64B*) (Morgante *et al*. 2015), while a wing morphology screen shared 16 genes (*bru1, Bx, CG1358, CG14926, Con, dally, dar1, Dgk, ds, Dys, Egfr, Lar, luna, pip, RhoGEF64C, Sp1*) (Pitchers *et al*. 2019). The overlap between these studies suggests that modifiers of robustness for flight performance also impact other traits, affirming their pleiotropic nature and their significance in quantitative genetic variation. These findings underscore the importance of studying the genetic architecture of robustness as it can impact the outcome of natural selection on quantitative traits.

## Conclusions

We present results from four analyses across four sex-based phenotypes surveying different facets of the genetic architecture of robustness for flight performance. Several of the variants were shared between sexes, though many more differed between them. Future studies in other phenotypes should consider evaluating both the mean and coefficient of variation for their focal phenotype to better understand modifiers affecting robustness in a specific complex trait, as well as robustness in complex traits more generally. Doing so would provide a better survey of the genetic modifiers of robustness as a phenotype and allow researchers to glean greater insight into the mechanisms of evolutionary change.

## Acknowledgements

We thank Faye Lemieux for assistance with fly husbandry, Jim A. Mossman for assistance collecting data, and anonymous reviewers for their feedback. This research was funded by National Institutes of Health grant GM067862 (to DMR).

## Data accessibility statement

All phenotype data required to run the outlined analyses are available in Table S1 or using the DGRP2 webserver (http://dgrp2.gnets.ncsu.edu/). Supplemental tables and files are available at: https://doi.org/10.7910/DVN/RSJNW5.

